# Neurocomputational underpinnings of expected surprise

**DOI:** 10.1101/501221

**Authors:** Françoise Lecaignard, Olivier Bertrand, Anne Caclin, Jérémie Mattout

**Author notes:** **Corresponding author:** Françoise Lecaignard, INSERM U1028 - CNRS UMR5292, Centre de Recherche en Neurosciences de Lyon, Equipe DYCOG, Centre Hospitalier Le Vinatier (Bât. 452), 95, Boulevard Pinel, 69500 Bron, France.

## Abstract

Predictive coding accounts of brain functions profoundly influence current approaches to perceptual synthesis. However, a fundamental paradox has emerged, that may be very relevant for understanding hallucinations, psychosis or cognitive inflexibility. This paradox is that in some situations surprise or prediction error related responses can decrease when predicted and yet, they can increase when we know they are predictable. This paradox is resolved by recognizing that brain responses reflect precision weighted prediction error. This then presses us to disambiguate the contributions of precision and prediction error in electrophysiology. We report, for the first time, an experimental paradigm that may be able to meet this challenge. We examined brain responses to unexpected and expected surprising sounds, assuming that the latter yield a smaller prediction error but much more amplified by a larger precision weight. Importantly, addressing this modulation requires the modelling of trial-by-trial variations of brain responses, that we reconstructed within a fronto-temporal network by combining EEG and MEG. Our results reveal an adaptive learning of surprise with larger integration of past (relevant) information in the context of expected surprises. Within the auditory hierarchy, this adaptation was found tied down to specific connections and reveals in particular and crucially precision encoding through neuronal excitability. Strikingly, these fine processes are automated as sound sequences were unattended. These findings directly speak to applications in psychiatry, where it has been suggested that a specifically impaired precision weighting is at the heart of several conditions such as schizophrenia and autism.

## Introduction

Brain responses to surprise are essential to understand how the brain adapts to a changing or uncertain environment. In perception research, the abundant literature dedicated to surprise related electrophysiological components has largely contributed to frame perception processes into regularity learning, independently of attention engagement. This important turn leverages on influential computational predictive brain theories (Dayan et al., 1995; Friston, 2012), with predictive coding algorithm in particular (Friston, 2005; Spratling, 2016). Under this view, evoked responses are treated as surprise or prediction errors indexing the discrepancy between predictions established through regularity learning and current sensations (Auksztulewicz and Friston, 2016; Heilbron and Chait, 2018; Lumaca et al., 2018; Schröger et al., 2015). These dynamic errors drive belief updating to ensure an on-going adaptation to changes. However, recent work points to a fundamental paradox that clearly deserves attention to further refine perceptual models (Auksztulewicz et al., 2017; Fitzgerald and Todd, 2020; Heilbron and Chait, 2018; Meyniel, 2020; Southwell et al., 2017; Walsh et al., 2020). Namely, brain activity increases with surprise, but it also increases when surprise becomes more predictable, in other words when surprise decreases.

Interestingly, predictive brain theories resolve this paradox by considering the precision (or confidence) the brain assigns to predictions and sensory inputs so that evoked responses to surprise would reflect *precision-weighted* prediction errors. Precision provides a formal way to control the gain of these errors according to their contextual relevance for an efficient and flexible perceptual processing (Clark, 2013; Friston, 2010; 2008a; Mathys et al., 2014). Remarkably, this complete and context-sensitive view highlights the fact that surprise processing therefore depends on the context, as well as adaptation to environmental changes. And the prediction error induced by a surprising event (a sensory input deviating from a regularity) is weighted differently depending on whether it is delivered within a structured (or predictable) environment or not, yielding expected and unexpected surprises, respectively. Predictability emerges from hierarchical statistical structure applying to every element of the sensory stream so that they each convey more information (about each hierarchical level rule) than the same elements delivered in a non-hierarchical structure. In perception as hierarchical regularity learning (or perceptual learning), higher-order belief updating modulates the precision of first-order (sensory) prediction errors. This implies at the sensory level: *i)* a higher precision afforded by predictability, yielding efficient belief updating, hence *ii*) lower prediction errors. This opposite effect of predictability on precision and prediction error makes this experimental manipulation a key to disambiguating their respective contribution to brain responses to further address perceptual processes and their (automatic) adaptation to changes.

In this study, we sought to shed light on this timely question by proposing an experimental paradigm that may be able to meet this challenge. Precisely, we manipulated the predictability of a binomial sound sequence in the aim to alter separately the contributions of precision and prediction error in auditory evoked responses. We conducted simultaneous EEG and MEG recordings (Lopes da Silva, 2013) to reveal related subtle changes of brain activity during passive listening. Trial-by-trial modelling of reconstructed cortical responses showed an automatic adaptation of learning to predictability (as expected under hierarchical learning), which here translates into a larger account of past (relevant) information under the more structured environment. Critically, we next examined the physiological counterpart of such adaptation using dynamic causal models (DCM) and it was measured in the synaptic connectivity in a fronto-temporal network which revealed separated accounts for precision weights and prediction errors. In addition to shedding light onto adaptive perceptual processes, these findings illustrate how predictive coding has the potential to serve perception research providing its key constructs are explicitly at play in the modelling of brain responses.

## Results

We report findings from simultaneous EEG-MEG recordings in 20 healthy volunteers undergoing a passive auditory oddball paradigm in two contexts of predictability: surprising sounds (frequency *deviants*, using the oddball terminology) were delivered either within a purely deterministic context (condition PC, predictable context), or a pseudo-random one (condition UC, unpredictable context) (Figure 1.A). Prior to addressing the automatic context-sensitivity of auditory processing, two control analyses were conducted to test the implicit learning of the two different statistical structures; it was demonstrated by a typical sensor-level evoked response analysis described in a previous report (Lecaignard et al., 2015) (Figure 1.B), and to test the perceptual learning at play for both contexts (PC, UC) within a fronto-temporal connectivity, in line with previous pioneering work in the field (Bayesian learning; Ostwald et al., 2012; Effective connectivity: Garrido et al. 2009). This second aspect is provided in the supplementary data. Here, using trial-by-trial computational modelling we tested whether the brain adapts its learning style to the contextual manipulation during unattentional listening. Results provided clear predictions regarding the mapping of precision and prediction error onto physiology, which we next tested using DCM (Bastos et al., 2012; Kiebel et al., 2009). A general view of this neuro-computational approach is provided in Figure 2.

**Figure 1.**
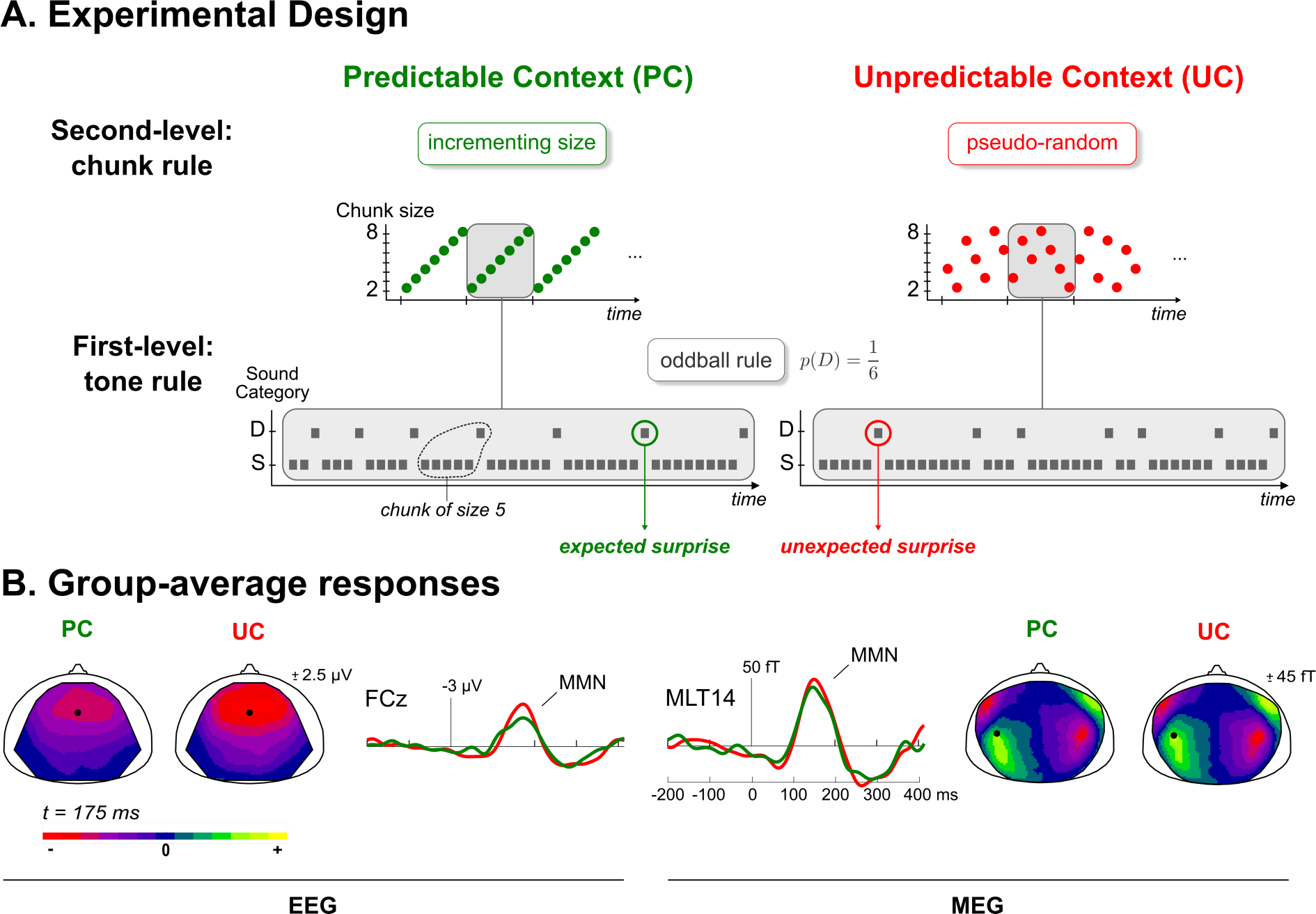
**A. Experimental Design.** Schematic view of the predictability manipulation (second level) applying to typical oddball sound sequences (first level). Predictable context (left, green) involves cycles of ordered transitions between segments of repeating standards (chunks), which become shuffled in the Unpredictable context (right, red). Deviant probability remains the same in both context (p = 1/6). Grey rectangles delineate an exemplary cycle for both sequences. S: Standard, D: Deviant. **B. Group-average difference responses**. For each modality (EEG, left; MEG, right), scalp maps of grand-average difference (deviant – standard) responses at the peak of the MMN (t=175 ms) for both contexts (PC: green; UC: red). Traces at sensors showing a significant MMN (EEG: FCz; MEG: MLT14) indicate the reduced MMN amplitude under predictability.

**Figure 2.**
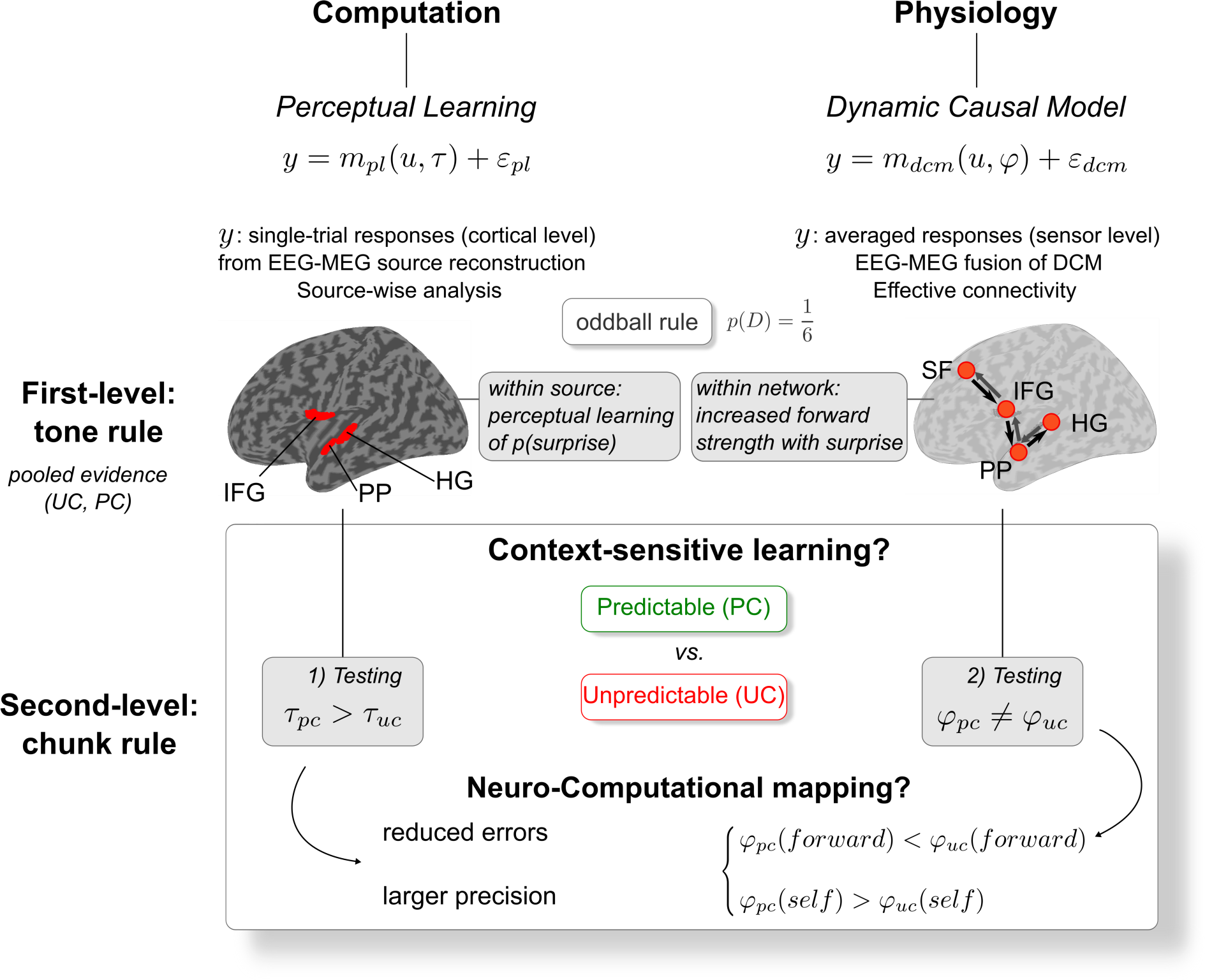
Neuro-computational framework. Simplified representation of the current approach deployed to address the automatic adaptive learning at play during auditory processing, and to disambiguate the mapping of precision-weighted prediction error onto physiology. Abbreviations: dynamic models: pl (perceptual learning), dcm (dynamic causal model); cortical sources: HG (Heschl’s gyrus), PP (planum polare), IFG (inferior frontal gyrus), SF (superior frontal); experimental contexts: PC/pc (predictable context), UC/uc (unpredictable context). D: deviant.

### Contextual adaptation of perceptual learning

This first analysis (Figure 2, lower-left panel) aims at testing the automatic adaptation of perceptual learning that would account for the reduced evoked response measured in the predictable context (Figure 1.B; Lecaignard et al., 2015). In a pre-analysis (Figure 2, upper-left panel), we controlled that the processing of sound - whatever the context - involves the perceptual learning of the oddball rule (see Supplementary data). We fitted dynamic learning models and static ones to single-trial cortical responses, with each model each furnishing a different cognitive account of the inter-trial variability, related to belief updating and change detection, respectively. Note that single-trial responses correspond to the reconstructed activity of six cortical sources at play within a fronto-temporal network using the fusion of EEG-MEG data (see Methods). They could be located bilaterally in Heschl’s gyrus (HG), the Planum Polare (PP) and the inferior frontal gyrus (IFG). Separate source-wise Bayesian model comparisons all revealed the outperformance of Bayesian learning model, especially at the latency of the MMN. Importantly for the current analysis, this winning model involves a temporal integration window to modulate the influence of past events and whose size is defined by a time constant parameter, *τ*. Beyond the memory-based interpretation (the brain may arguably not be able to deal with long-gone, past information), this parameter endows the learning model with a flexible way to integrate past information and formalizes brain adaptation to its environment. From supplemental Figure S1, it can be seen that the precision-weighted prediction error, here defined as a Bayesian Surprise (see Methods), decreases with *τ*, reflecting the better predictions induced from a larger account of past information. Besides, simulated MMNs for a given *τ* value show a similar amplitude across contexts (Figure 3.A). This suggests that the reduced MMN measured in condition PC does not emerge from the sound sequence structure alone. This pleads for a hierarchical processing enabling an adaptive learning: we expect a larger *τ* under the more predictable sequence. We performed model inversion at every time sample exhibiting statistical significance in the pre-analysis. For each cortical source and each parameter, posterior estimates were averaged across samples; they were subsequently compared by a repeated-measures ANOVA across factors *Condition* (UC, PC), *Hemisphere* (Left, Right) and Sources (HG, PP, IFG). Larger *τ* values were found with condition PC compared to condition UC (F_(1,19)_ = 10.13; p = 0.002). On average across sources, *τ* was equal to 16.0 and to 21.3 with UC and PC, respectively (Figure 3.B). The ANOVA showed no other significant effect (all p>0.12). As can be seen in Figure 3.C, the different *τ* values in UC and PC generate different down-weightings of past observations, with a larger amount of information integrated during perceptual learning in the predictable context.

**Figure 3.**
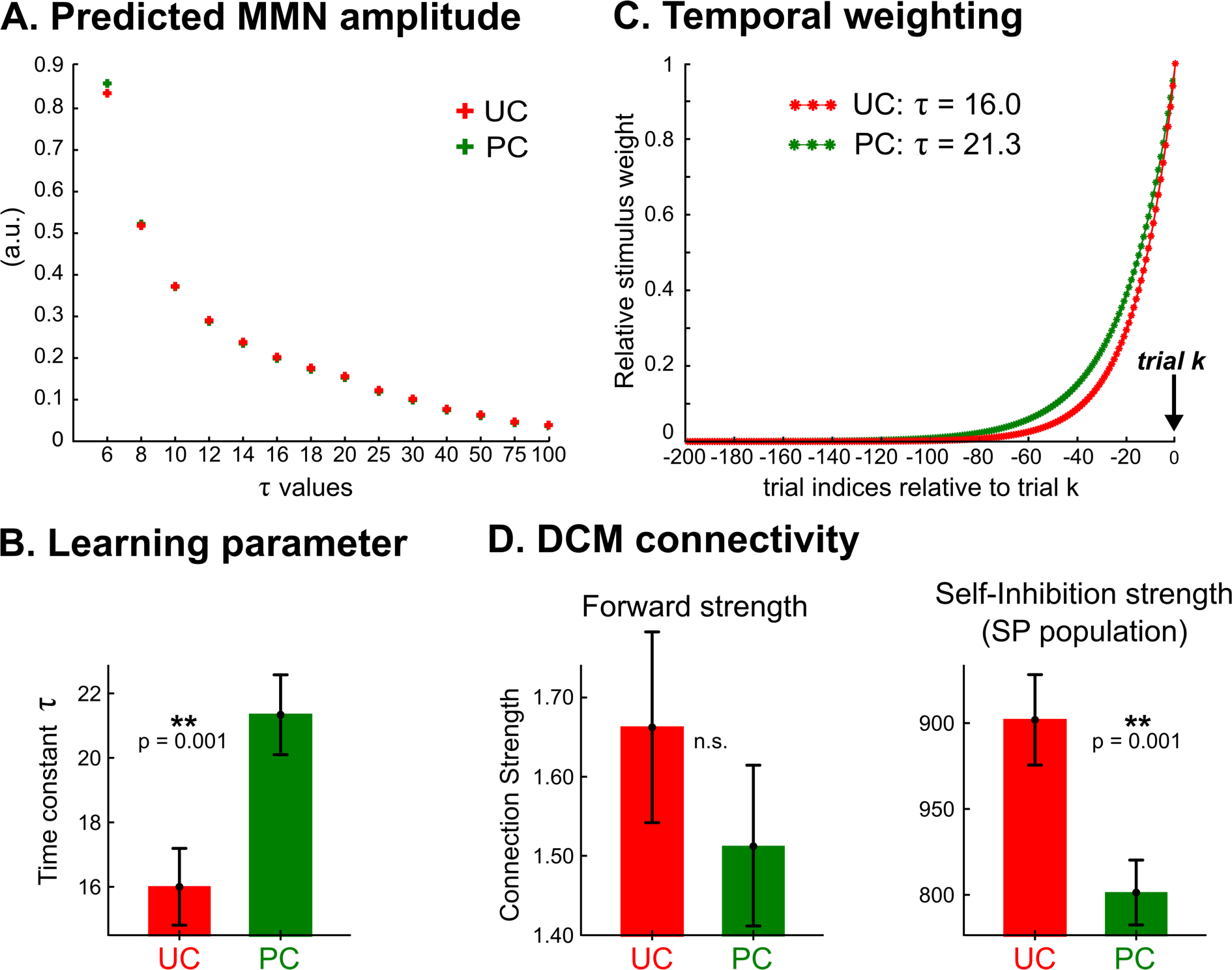
Effect of predictability on auditory processing. Panels A, B and C refer to the cognitive modelling (perceptual learning), and panel D to its physiological counterpart (DCM). A) Learning model predictions of the MMN amplitude (group average) as a function of *τ* for UC (red) and PC (green) sound sequences (see Methods). Note that in both contexts, MMN amplitude decreases similarly as *τ* increases. B) Effect of predictability on learning parameter *τ*. Posterior estimates of *τ* averaged at the group-level and over the six clusters exhibited a significant difference between conditions. C) Downweighting of past observations obtained with the posterior estimates of *τ* for conditions UC (red) and PC (green), respectively. D) Effect of predictability onto effective connectivity obtained with the fusion of EEG and MEG DCMs. Posterior estimates of forward (left) and self-inhibition (right) strengths measured in both conditions (UC: red, PC: green); averaged over the group and over the connections within the DCM. SP: superficial pyramidal; a. u.: arbitrary units.

### Separate neural correlates of prediction error and precision

We next examined how this adaptation of learning manifests at the physiological level, by measuring the predictability effect onto the effective connectivity at play during oddball processing (Figure 2, lower-right panel). Such connectivity was previously inferred from averaged responses evoked by surprising (*deviant*) and regular sounds (*standard* preceding a deviant) in the two contexts using dynamic causal models (Figure 2, upper-right panel). As described in the supplementary data (Figure A3), Bayesian model comparison at the group level was in favor of a four-level hierarchy deploying over a fronto-temporal network with bilateral stations in HG, PP and IFG, and in an additional superior frontal contribution (SF). The winning DCM comprises intra-hemispherical reciprocal synaptic connections and subcortical inputs in bilateral primary auditory area (HG) and the inferior frontal gyrus (IFG). Importantly, the modulation of synaptic connectivity induced by the change of input (from a standard to a deviant) strongly supports the neural implementation of predictive coding message passing along the cortical auditory hierarchy. In the present analysis, and like in the previous trial-by-trial modelling one, this winning model was next inverted separately for UC and PC responses to re-evaluate specific model parameters, namely the intrinsic and extrinsic connection strength as well as their modulation by input change, for which we expect to disclose the putative contextual adaptation effect (Figure 2, lower-right panel). Precisely, based on the reported adaptation of learning to predictability (larger *τ*), we expect:

1. Lower forward connection strength in condition PC, if it was to encode the (precision-weighted) prediction error passing. These errors are effectively reduced when the temporal integration window is lengthened.
2. Lower self-inhibition connection strength in condition PC, if it was to encode precision as suggested by previous work (Auksztulewicz et al., 2017; Brown and Friston, 2013; Fogelson et al., 2014; Moran et al., 2013). This intrinsic DCM parameter controls the excitability of superficial pyramidal cells whose cognitive role could be related to the computation of prediction errors (Bastos et al., 2012; Feldman and Friston, 2010; Friston, 2008a). As already mentioned, structured contexts yield precision increase which amplifies prediction errors according to their larger informational value.

Our inversion scheme was here again informed by both EEG and MEG modalities (we here refer to their posterior fusion, *p-MEEG;* see Methods). Resulting subject-wise DCM parameter estimates were computed using Bayesian model averaging (BMA; Penny et al., 2006) in the p-MEEG modality for each condition.

To test first prediction, we conducted a repeated-measures ANOVA over forward connection strength estimates, with factors *Condition* (UC, PC), *Hemisphere* (Left, Right) and *Level* (Temporal, Fronto-Temporal, Frontal). As can be seen in Figure 3.D (left), predictability yielded a reduced forward connection strength on average across the network (average with standard error: 1.51 ± 0.10 and 1.66 ± 0.12, in PC and UC resp.), an effect which was largest at the lowest level of the network (PC / UC. Level Temporal: 1.52 ± 0.10 / 2.05 ± 0.24; Level Fronto-Temporal: 1.41 ± 0.16 / 1.50 ± 0.21; Level Frontal: 1.61 ± 0.24 / 1.44 ± 0.17). However, these effects did not reach statistical significance (*Condition*: F_(1,19)_=0.84; p=0.36; *Condition×Level*: F_(1,19)_=1.82; p=0.17). The second prediction was tested with a similar ANOVA over BMA self-inhibition estimates, here factor *Level* comprises the four bilateral sources of DCM architecture (HG, PP, IFG, SF). As expected, significantly reduced self-inhibition connection strength was measured in condition PC compared to condition UC (801.4 ± 18.9 and 901.9 ± 26.3, resp.; F_(1,19)_ = 10.52; p = 0.001), as shown in Figure 3.D (right). This result establishes a link between self-inhibition and precision, as both adapt consistently to predictability (Figure 2, lower panel). Finally, the ANOVA was conducted on standard-to-deviant modulation applying on forward connection strength and self-inhibition parameters. We found no significant main effect of *Condition* (forward: F_(1,19)_=2.50; p=0.12; self-inhibition: F_(1,19)_=0.01; p=0.90). These latter results suggest that the contextual effect of our predictability manipulation apply indistinctively to standard and deviant stimuli processing.

## Discussion

This work addressed the hierarchical processing of sensory information during unattentional listening through its adaptation to the statistical structure of the environment. Complementary cognitive and neurophysiological modelling of oddball responses collected during different contexts of predictability reveals the occurrence of perceptual learning within a fronto-temporal hierarchy, as expected in the predictive coding framework. Moreover, it formally demonstrates that this predictive process is shaped by predictability in a way that optimizes the integration of relevant sensory information over time. Computationally speaking, this adaptation relies on the tuning of the precision weighting of prediction errors, a process which is known to be a cornerstone of Bayesian information processing (Mathys et al., 2014). We therefore show with a mechanistic and dynamical approach that during passive oddball listening, the more structured is the environment, the more efficient is sound processing. Besides, for the first time the neural encoding of precision weight could be distinguished from the prediction error *per se* and directly attributed to inhibitory mechanisms. Remarkably, the hypothesis of evoked responses reflecting hidden precision-weighted prediction errors (Friston, 2005) that was considered along this work is substantiated empirically by the present findings obtained with complementary neuro- and computational generative models of evoked responses.

Rare but robust empirical supports for predictive coding at play during oddball processing have been reported this past decade, obtained at the psychological level with trial-by-trial computational modelling (Lieder et al., 2013; Ostwald et al., 2012; Stefanics et al., 2018; Weber et al., 2020) and at the physiological level with DCM (Chennu et al., 2016; Fogelson et al., 2014; Garrido et al., 2009; Lumaca et al., 2021; Moran et al., 2013). Novel evidence at the both levels of analysis is provided here in the controlled work using exactly the same brain data informed by simultaneous EEG and MEG. The major contribution of this work, in the perspective of testing predictive coding more finely, lies in the fact that we could evidence the automatic tuning of the precision weighting of sensory errors and relate it to inhibitory mechanisms. This was made possible with the proposed model-driven contextual manipulation, where a second-order rule applies on the first-order oddball one. At the cognitive level, our results demonstrate quantitatively brain’s ability to grasp implicitly the larger informational content of the predictable context to derive a more informed and more efficient learning at the sensory level (by means of a larger temporal integration window). The larger efficiency in sensory processing translates into more accurate sensory predictions and more rapid adaptation to unexpectedness (sensory precision is related to the learning rate) (Mathys et al., 2014). This could provide a mechanistic explanation to the better task performances reported in target detection during the listening of regular auditory streams akin random ones (Southwell et al., 2017), and also during the processing of words and music stimuli under contextual expectancy (Tillmann et al., 1998). Similar facilitation of sensory processing in a structured environment was also suggested in a visual discrimination study (Rohenkohl et al., 2012), where temporal expectation was found to decrease reaction time, and could be associated to an increase of the sensory gain using a diffusion model. This result is comparable to the present precision adaptation, that here occurs without attentional (active) processing. At the neural level, our findings support the view of forward connections carrying precision-weighted prediction errors computed in supragranular cortical layers, and establish empirically a direct link between precision and self-inhibition within this layer. Several recent DCM studies strongly supported this mapping, as they reported consistent modulations of self-inhibition by some experimental manipulations hypothesized to influence precision under predictive coding, namely a cholinergic neuromodulation (Moran et al., 2013), sensory precision (Brown and Friston, 2012), selective attention (Brown and Friston, 2013), and predictability (Auksztulewicz et al., 2017). Here, it is the automatic contextual adaptation of sensory processing that we reveal *computationally* and *physiologically* that fills the missing link enabling to relate precision and self-inhibition directly. From a methodological perspective, DCM findings add to recent efforts to increase model plausibility to account for electrophysiological data (Phillips et al., 2016; Pinotsis et al., 2017).

Note that the study does not address the computational role of each DCM level. Rather we considered a single hierarchical-level learning based on a simple Bernoulli distribution (other, slightly more complex transition probability models could be considered, e.g. see Meyniel et al., (2016)) because it was equipped to reveal an adaptive learning in combination with the contextual manipulation. Such adaptation requires hierarchical learning to emerge, as higher levels compute errors and related precisions that pertain to more stable or slow-changing (contextual) stimulus features (Friston and Kiebel, 2009). In addition, increasing second-order statistic reliability has been theoretically demonstrated to increase the precision-weighting of sensory errors (Friston, 2008b; Kanai et al., 2015). The present report of automatic adaptation of perceptual learning and the related tuning of underlying connectivity fits very well with the top-down influence of contextual learning on sensory processing.

Future research in testing predictive brain theories for perception has to address the mechanisms which subsume the adaptation of the learning process that we observed, including the dynamics of precision tuning. From a psychological perspective we aim at investigating more finely the present predictability adaptation in relation to attentional processes, guided by the computational account of attention (Friston, 2005). Under this view, attention serves to collect contextually-informative sensations to optimize perception and learning (Auksztulewicz and Friston, 2016; Friston, 2005; Parr and Friston, 2018), through either the precision weighting of sensory channels (Feldman and Friston, 2010) or action. The view of attention as the tuning of precision echoes our findings obtained without participant’s awareness of the experimental manipulation. Interestingly, predictive brain theories have led to consider under the same framework the opposite effect of voluntary attention (an increase) and predictability (a decrease) on evoked response amplitudes reflecting precision-weighted prediction errors. In Chennu et al. (2013) and in Auksztulewicz and Friston (2015), both factors could be manipulated orthogonally using different task instructions during perception of oddball-like sound sequences. These two studies revealed different modulations of mismatch responses at different latencies, and related attention to self-inhibition using DCM, respectively. Chennu and colleagues reported a reduced MMN when attention was explicitly engaged towards (local) tone transitions compared to (global) multi-tone patterns. This fits with the present reduced MMN in the predictable condition considering that a more informed learning of the oddball rule (providing better predictions) could be at play either through an explicit attentional engagement or, in our case, through the implicit learning of the contextual information. Therefore, we argue that likewise but without the voluntary orientation of attention, predictability acts as an implicit attentional process, enhancing the efficiency of sensory processing. Similar effect of voluntary attention and predictability on precision (an increase) emphasizes the great potential of separating prediction error and precision accounts in order to predict (and test) their respective effects on evoked responses, instead of addressing precision-weighted prediction errors as a whole (for review Heilbron and Chait, 2018). Bridging passive predictability processing and voluntary attention opens the way to mechanistically investigate attentional capture. This would involve experimental manipulations of precision and prediction error, and appropriate hierarchical dynamic models to assess underlying activity, and feasibility was demonstrated here. We expect in particular novel insights from the manipulation of precision (in addition to typical modulations of global precision-weighted prediction errors), to assess whether voluntary attention can emerge from specific precision evolution reflecting a form of evidence accumulation.

## Conclusion

A contextual manipulation of oddball paradigm was used to disentangle prediction error and precision neural representations. Contextual effect was found to increase the extent of temporal integration of past information, which implies lower sensory prediction errors amplified by a larger precision weighting. Findings in this paper 1) demonstrate the conclusive power of modelling approaches combining neuronal and cognitive levels and 2) emphasize the importance of accounting for the encoding of precision weighting when investigating perceptual learning and decision-making. Unfolding the mechanisms of precision tuning and encoding, especially at an implicit level, is a potentially critical step for clinical applications as alterations of these processes have been suggested to be at the core of several psychiatric disorders (Adams et al., 2013; Friston, 2020; Haarsma et al., 2020; Lawson et al., 2017). Applying such a simple oddball paradigm, only involving passive listening, coupled with computational and neurophysiological modelling could be of great value in this context.

## Methods

### Participants

Twenty healthy volunteers (10 female, mean age 25±5 years, ranging from 18 to 35) participated in the study. All participants were free from neurological or psychiatric disorder, and reported normal hearing. All participants gave written informed consent and were paid for their participation. Ethical approval was obtained from the appropriate regional ethics committee on Human Research (CPP Sud-Est IV - 2010-A00301-38).

### Experimental design

Predictable and unpredictable sound sequences embedding a typical frequency oddball rule (conditions PC and UC, respectively) were used in the present study. Participants were instructed to ignore the sounds and watch a silent movie of their choice with subtitles. Predictable sound sequences comprised 16 cycles that were each made of a repeating 42-tone pattern following the deterministic incrementing rule depicted in Figure 1.A. Unpredictable sequences corresponded to pseudo-random oddball sequences typically used in oddball paradigms, with specific controls for the number of standards in between two deviants. Despite their differing statistical structure, both sequence types had the same deviant probability (*p* = 0.17) and the same distribution of deviants among standards (there were exactly the same number of chunks of repeating standards before a deviant in both conditions, with chunk size varying from 2 to 8 standards). Further details about stimuli and sequences can be found in Lecaignard et al. (2015). All stimuli were delivered using Presentation software (Neurobehavioral Systems, Albany, CA, USA).

### Data acquisition and pre-processing

Simultaneous MEG and EEG recordings were carried out in a magnetically shielded room with a whole-head 275-channel gradiometer (CTF-275 by VSM Medtech Inc.) and the CTF-supplied EEG recording system (63 electrodes), respectively. Signal was amplified, band-pass filtered (0.016–150 Hz), digitized (sampling frequency 600 Hz) and stored for off-line analysis. First-order spatial gradient noise cancellation was applied to MEG signal. EEG reference and ground electrodes were placed on the tip of the nose and left shoulder, respectively. Digitization of electrode locations (Fastrak, Polhemus, Colchester, VT, USA) and acquisition of individual T1-weighted magnetic resonance imaging images (MRIs, Magnetom Sonata 1.5 T, Siemens, Erlangen, Germany) were conducted for coregistration purposes in distributed source reconstruction (see below).

Pre-processing of data using the ELAN package (Aguera et al., 2011) and Matlab routines included the following: rejection of data segments affected by head movements (larger than 15 mm relative to the average position over sessions) or SQUID jumps, power-line filtering (stop-band filters centered on 50 Hz, 100 Hz, and 150 Hz with bandwidth of ± 2 Hz), independent component analysis (ICA) correction for ocular artifacts (EEGlab routines, http://sccn.ucsd.edu/eeglab/index.html), rejection of trial epochs (from −200 ms to 410 ms after stimulus onset) with signal amplitude range (over entire epoch) exceeding 2000 fT for MEG data and 150 μV for EEG data, and 2-45 Hz band-pass digital filtering (bidirectional Butterworth, 4th order). Importantly, we only used time epochs that survived the procedures applied for artifact rejection for both modalities. Finally, trial epochs were imported in SPM (Wellcome Department of Imaging Neuroscience, http://www.fil.ion.ucl.ac.uk/spm) and were down-sampled (200 Hz) for data reduction and low-pass filtered (20 Hz low-pass digital filter, bidirectional Butterworth, 5th order). Group-average deviance responses obtained in EEG and MEG in both UC and PC conditions are shown in Figure 1.B.

### EEG and MEG evoked responses

Averaged responses evoked by standards just preceding a deviant and by deviants were considered for all DCM analyses. EEG responses were re-referenced to the averaged mastoid electrodes. Time interval of 0 ms to 220 ms after sound onset was used for model inversion. It was defined from sensor-level (EEG and MEG) statistical analysis on deviance responses to ensure it encompasses the MMN (and no later components). A Hanning window was applied to time-series to ensure that system’s dynamics was set to zero before being excited. Data reduction achieved within the inversion procedure led to the selection of 8 (±2.7) and 13 spatial modes with EEG and MEG data, respectively (on average across subjects).

### Trial-wise reconstructed cortical data

Learning model were confronted to trial-by-trial timeseries collected with EEG and MEG. Single-trial data was obtained in a preparatory step involving the distributed source reconstruction of EEG-MEG data. Advanced methods were employed for source inversion with realistic forward models for both modalities(Gramfort et al., 2010), Bayesian framework enabling Multiple Sparse Priors (Friston et al., 2008; Mattout et al., 2006), EEG-MEG fusion (Henson et al., 2009), and group-level inference (Litvak and Friston, 2008). Source inversions were all performed with the SPM software (SPM8 release). First, in a prior study (Lecaignard et al., n.d.) six cortical clusters could be identified from the inversions of early and late (MMN) mismatch peaks. These sources were located in the left and right Heschl’s gyrus (HG), Planum Polare (PP) and inferior frontal gyrus (IFG) (Supplementary Figure A1). Critically, they subsequently served as spatial priors to constrain the distributed inversion of entire single-trial epochs (from −200 ms to +410 ms, 674 trials per condition per participant). Within each cluster and for each trial, reconstructed cluster-node activities were averaged to derive a cluster-level and single-trial trace being informed by both EEG and MEG data.

### Computational modeling (cognitive models)

We considered the learning model with highest evidence in the control pre-analysis (see Supplementary Data). It assumes that the brain learns from each stimulus exposure the probability *μ* to have a deviant, in order to predict the next sound category *u* (with *u* = 1 in the case of a deviant and *u* = 0 in the case of a standard). We define *u* ∼ *Bern* (*μ*) with *Bern* the Bernoulli distribution, and *μ* ∼ *Beta* (α, β) with α and β the parameters of the distribution *Beta*, corresponding in the current case to deviant and standard counts, respectively. At trial *k*, we have:

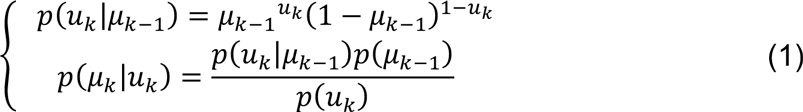

The posterior distribution of *μ* is in the form of a *Beta* distribution (*Beta* distribution is conjugate to the *Bern* distribution), leading to the following updated expression of *μ* at trial *k*:

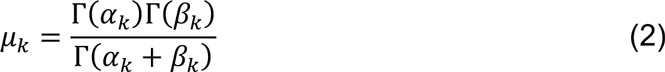

With Γ the Gamma Euler function, and α and β following update equations:

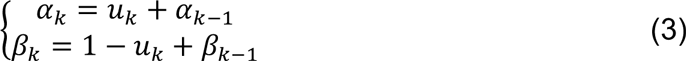

Precision-weighted prediction error is the Kullback-Leibler (KL) divergence between the prior and the posterior *Beta* distributions of *μ*, also referred to as a Bayesian Surprise (BS) in the MMN study by Ostwald et al. (2012). At trial *k*, it expresses as:

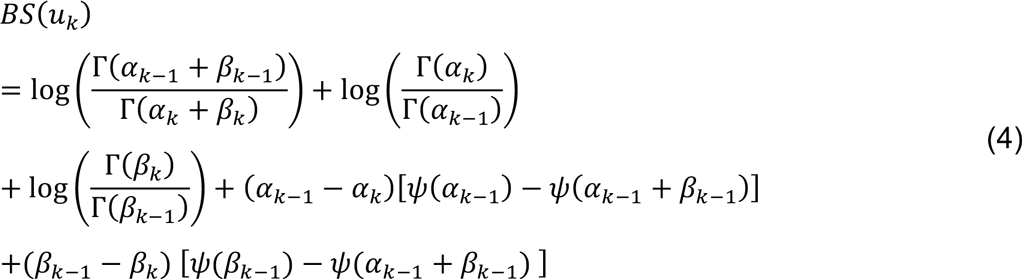

With *ψ* the digamma Euler function. Importantly for our investigation, the size of the temporal integration window was parameterized by *τ* which enters standard and deviant count updates as follows:

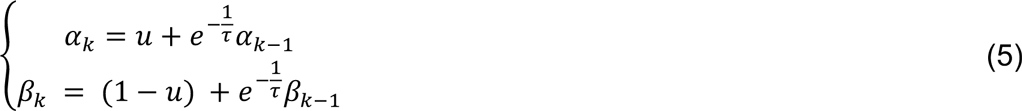

From equation Eq. 5 we see that the larger the *τ*, the larger the weight applying to past observations, leading to a more informed learning (an illustration can be found in Figure 3.C). Variation of BS with the size of the temporal integration window is shown in supplemental Figure S1.

#### • Group-level simulated MMN

In order to test if sound transition alone, which differ across contexts, is sufficient to explain the reduced MMN observed under predictability, we simulated group-level MMN for different values of *τ*. For each subject, we computed individual trial-by-trial precision-weighted prediction error trajectories induced by UC and PC sequences for *τ* in {6,8,10,12,14,16,18,20,25,30,40,50,75,100}, using VBA (Daunizeau et al., 2014). We then selected the values obtained for each standard preceding a deviant and for each deviant (in keeping with sensor-level trial rejection). Using exactly the same procedure that had been used to compute the event-related difference response at the sensor level, we could simulate the group-average MMN amplitude in conditions UC and PC (arbitrary units). Results are provided in Figure 3.A.

#### • Model inversion

Using the VBA toolbox (Daunizeau et al., 2014), inversion of the learning model was performed separately in each context, in each cluster, and at every time sample of the peri-stimulus period that exhibited this model as winning in the pre-analysis (the number of samples consequently varied across clusters, leading to a total of 33 samples). Bad trials (with regard to sensor-level artifact rejection) were processed in a way that corresponding signals would not corrupt parameter optimization (note that related stimuli still entered learning dynamics because they were observed by the brain). Detailed model specifications in VBA are provided in the Supplementary data. Overall, 66 inversions were conducted per subject (1 model, 2 conditions, 33 samples). Each inversion provided a posterior estimate for parameter *τ* informed by EEG and MEG data.

#### • Statistical analysis

We assumed a constant value of *τ* within the time interval used for model inversion (spanning the MMN). We therefore averaged *τ* estimates across samples for each cluster. Predictability effect could thus be analyzed by conducting a repeated-measures ANOVA on these posterior estimates with factors *Condition* (UC, PC), *Hemisphere* (Left, Right) and *Sources* (HG, PP, IFG).

### Neural modeling (Dynamic Causal Modeling, DCM)

DCM architecture was inferred in a previous control analysis (Supplementary Data). It involves four hierarchical levels defined from the sources of the early and late (MMN) components that we previously reconstructed using fused EEG-MEG inversion (Lecaignard et al., 2021). In addition to the 6 bilateral clusters identified over Heschl’s Gyrus (HG), the Planum Polare (PP), and the Inferior Frontal Gyrus (IFG), a bilateral superior frontal level (SF) was needed to increase model accuracy. DCM inputs were found in HG and IFG stations.

#### • Model space

We looked for predictability effect on the synaptic connectivity established for deviance processing. We thus considered the same model space made of 14 models (fully described in the Supplementary Data) here confronted against responses in condition UC and PC separately.

#### • Model inversion

DCM analyses were performed with SPM12 (Wellcome Department of Imaging Neuroscience, http://www.fil.ion.ucl.ac.ik/spm). We used the canonical micro-circuit (CMC) neural mass model (Bastos et al., 2012; Brown and Friston, 2013) to exploit its relevance to test predictive coding predictions. We used default values of SPM12 as prior expectations and prior variance for each DCM parameter to be estimated. Each DCM inversion involved standard response as the initial state of the system and deviant response resulting from the experimental perturbation. Regarding DCM sources, we maintained dipole locations fixed (but not their orientation) to inform spatially DCM inversion with the group-level MEG-EEG information. For each modality (EEG, MEG), the forward model used for DCM inversion was an advanced realistic Boundary Element Model (BEM) computed with Openmeeg software (Gramfort et al., 2010).

#### • Fusion of EEG and MEG DCMs

To date, the fusion of EEG and MEG data within the generative model of DCM is not implemented. Our approach consisted in performing EEG and MEG DCM inversions separately (using exactly the same parameter priors), and in a second step, exploiting the Bayesian framework of DCM to derive posterior estimates informed by both modalities (referred to as p-MEEG). Precisely, the proposed procedure rests on the assumption of conditional independence of EEG and MEG data under the quasi-static approximation of Maxwell equations, which is largely admitted for signals below 1kHz (as is the case here). Denoting EEG and MEG data by *y_EEG_* and *y_MEG_*, respectively, we have:

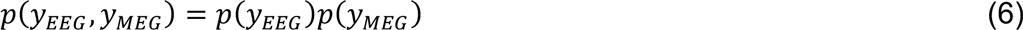

And posterior model evidence of model *m* can be approximated using unimodal EEG and MEG model evidences:

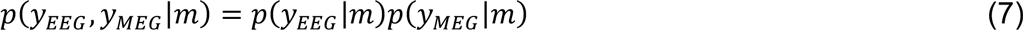

Consequently, ℱ*_p-MEEG_* the variational free energy approximation to p-MEEG model log-evidence could be obtained by:

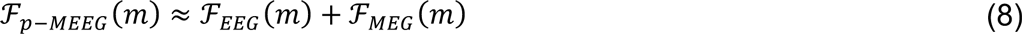

With ℱ*_EEG_* and ℱ*_MEG_* the free energy values for EEG and MEG respectively. Besides, the posterior distribution of some DCM parameter *θ* under model *m* writes given:

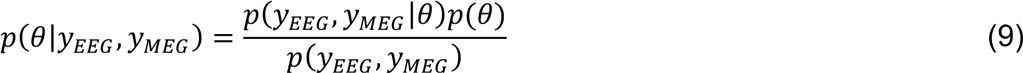

Which can be re-formulated as follows to reveal the posterior distributions of *θ* deriving from unimodal inversion of EEG and MEG data:

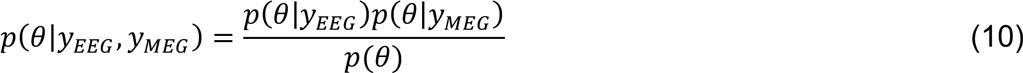

DCM approach assumes every parameter *θ* to have of the form of a Gaussian distribution. Hence prior distribution expresses as *q*(*θ*) ∼ 𝒩(*μ*_o_, *σ*_o_). We also denote *q*(*θ*, *y_EEG_*) ∼ 𝒩(*μ*_e_, *σ_e_*), *q*(*θ*, *y_MEG_*) ∼ 𝒩(*μ_m_*, *σ_m_*) and *q*(*θ*, *y_EEG_*, *y_MEG_*) ∼ 𝒩(*μ_p_*, *σ_p_*) the posterior distribution of *θ* given EEG data, MEG data and EEG-and-MEG data respectively. We have *μ*cd and *σ*cd the mean and variance of the distribution resulting from the multiplication of *q*(*θ*, *y_EEG_*) and *q*(*θ*, *y_MEG_*). From equation Eq.10 and the analytical expressions of *μ_em_* and **σ*_em_* (detailed in most statistic books), we derive:

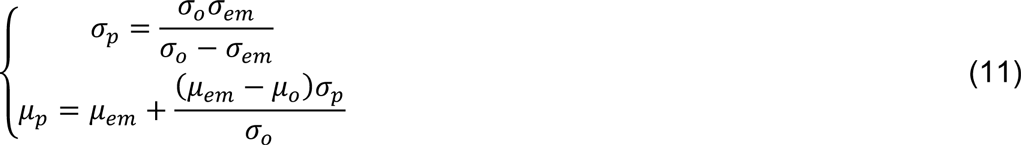

#### • Statistical analysis

We computed condition-wise individual BMA posterior estimates (in the p-MEEG modality) over model space. The analysis rests on four connections in particular: forward and self-inhibition connection strengths, and their respective trial-specific change (standard-to-deviant modulation). Note that DCM with CMC assumes two types of forward connections: a predominant one originating in superficial pyramidal (SP) subpopulation and targeting the spiny stellate (SS) cells of the higher-level source, and a secondary one linking subpopulation SP to higher-level deep pyramidal (DP) subpopulation (supplemental Figure S2, left). Here, we report only the effect of predictability on the former type. For the forward related parameters, we conducted a repeated-measures ANOVA over individual BMA estimates with factors *Condition* (UC, PC), *Hemisphere* (Left, Right) and *Level* (Temporal, Fronto-Temporal, Frontal); in the self-inhibition related parameter, this latter factor was replaced by factor *Source* (HG, PP, IFG, SF).

Throughout the paper, ANOVA analyses were performed using R software (The R Foundation, https://www.r-project.org/).

## Acknowledgments

We thank Sébastien Daligault and Pascal Calvat for programming support in interfacing with the CC IN2P3. We acknowledge CC-IN2P3 for providing computing resources and services needed for this work. We thank Emmanuel Maby and Gaëtan Sanchez for helpful discussions. Thank you to Karl Friston and Florent Meyniel for valuable feedback on the manuscript. This work was supported by a grant from the Agence Nationale de la Recherche of the French Ministry of Research ANR-11-BSH2-001-01 to AC and FL and a grant from the Fondation pour la Recherche Médicale (FRM) to OB and JM. This work was conducted in the framework of the LabEx CeLyA (“Centre Lyonnais d’Acoustique”, ANR-10-LABX-0060) and of the LabEx Cortex (“Construction, Function and Cognitive Function and Rehabilitation of the Cortex”, ANR-10-LABX-0042) of Université de Lyon, within the program “Investissements d’avenir” (ANR-11-IDEX-0007) operated by the French National Research Agency (ANR).

## Author Contributions

Conceptualization, AC, FL and JM; Methodology, FL and JM; Software, FL and JM; Investigation, AC and FL; Writing, AC, FL and JM; Funding Acquisition, OB, AC and JM; Supervision, OB, AC and JM.

## Declaration of Interest

The authors declare that they have no conflict of interests.

## Supplementary Data

Prior to testing the predictability effect onto perceptual learning and associated effective connectivity, we first conducted two preliminary analyses aiming at characterizing the processing of unexpected sounds (across both contexts) at the cognitive level (using trial-by-trial computational modelling) and the physiological level (using DCM). Findings in each analysis (but obtained with exactly the same data) reinforce pioneering work in the field (Garrido et al., 2009a; Ostwald et al., 2012) and evidence Bayesian learning at play in a fronto-temporal hierarchy.

Brain responses used for each modelling work (perceptual learning and DCM) are described in the main text (Methods section). Cortical clusters were identified at the latency of the MMN using fused EEG-MEG source reconstruction (Lecaignard et al., 2021). They concern bilateral sources in the Heschl’s Gyrus (HG), the Planum Polare (PP) and the Inferior Frontal Gyrus (IFG) (Figure A1.A). In addition, a bilateral contribution in the superior frontal (SF) lobe was introduced in DCM.

**Figure A1.**
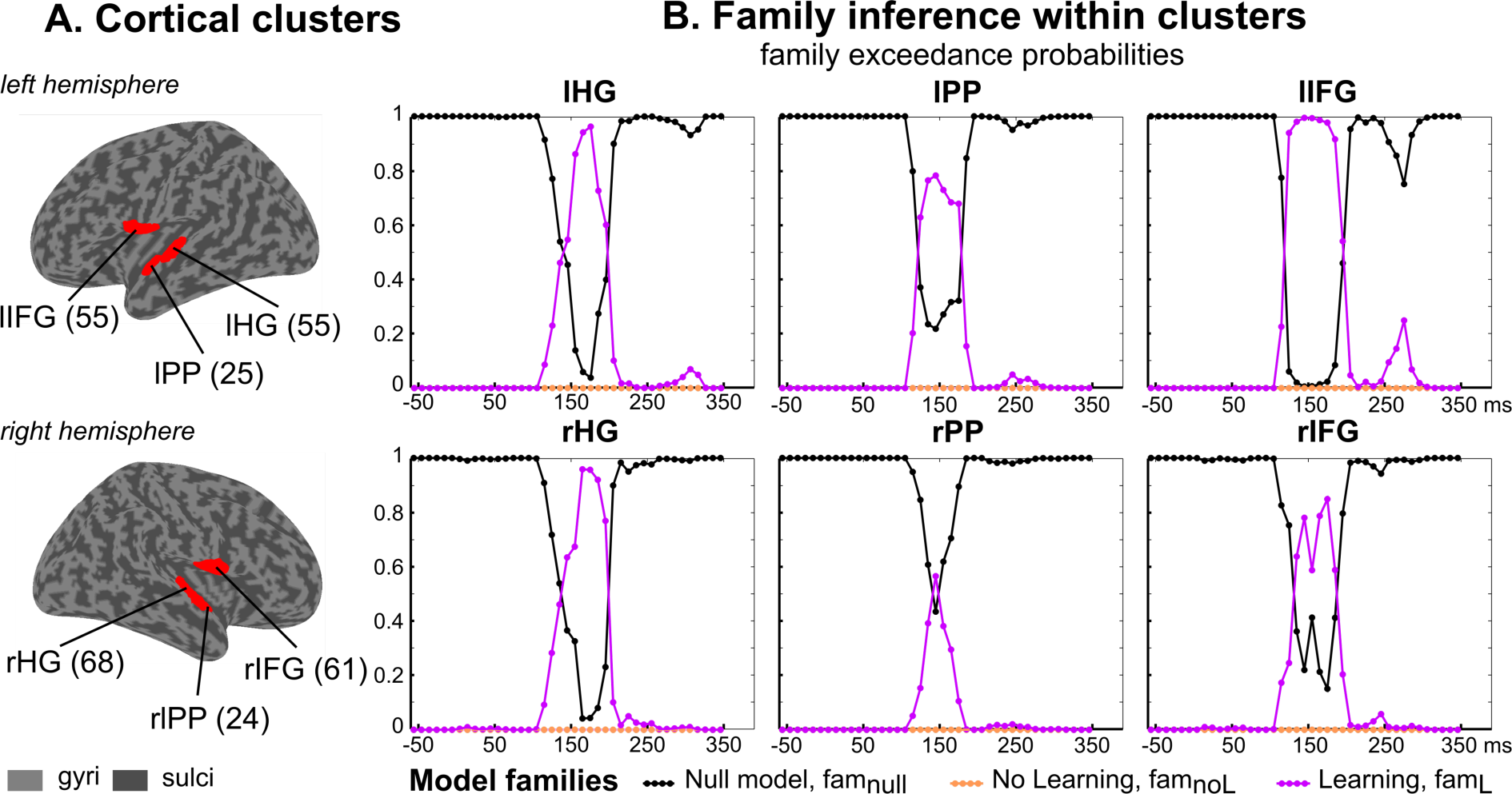
Computational modelling of deviance processing. A) Cortical clusters of interest are represented (red) over the inflated cortical surface of the SPM template brain (Mattout et al., 2007). These six clusters are left, right Heschl’s Gyrus (lHG, rHG), left, right Planum Polare (lPP, rPP) and left, right Inferior Frontal Gyrus (lIFG, rIFG). Total number of nodes in each cluster is indicated in parenthesis). Within each cluster, reconstructed single-trial traces (informed by EEG and MEG) were averaged over nodes and confronted to trial-by-trial model predictions at each time sample within [-50 +350] ms. B) Family-wise Bayesian model comparison. For each cluster and at each time sample, family inference provides the estimated posterior family exceedance probability of each model family (fam_null_: black, fam_noL_: orange, fam_L_: pink).

### A1. Perceptual learning of environmental regularities

Under predictive coding mismatch responses (including the Mismatch Negativity, MMN) elicited by unexpected deviants reflect prediction errors of a continuously updated model of the acoustic environment (Friston, 2005). We tested this hypothesis against alternative cognitive processes that did not involve perceptual learning (presented below).

#### 1. Methods

Models were all defined as a two-level linear model of the form:

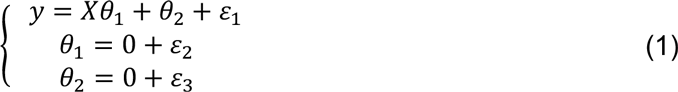

Where y indicates the reconstructed cortical activity informed by a fused EEG-MEG source inversion in the form of a vector of trial-by-trial activity at a particular sample of the peristimulus time; *X* is defined for each model and represents the predicted trajectory of precision-weighted prediction error over the sound sequence; {*θ*_1_, *θ*_2_) } and {*ε*_1_, *ε*_2_, *ε*_3_} refer to Gaussian observation parameters and Gaussian noise, respectively. Vector y was defined for each time sample of the [-50 +350] ms epoch and for each cluster of the MMN cortical network identified at the group level (6 clusters). We considered a model space of seven cognitive models partitioned into three families, largely inspired by models used in a previous tactile oddball study (Ostwald et al., 2012):

- Family *fam_null_* is made of a single model, the null model (M0) assuming that the brain response to every tone is the same, or equivalently that EEG fluctuations only reflect random noise (i.e. *θ*’ = 0 in Eq.1).
- Family *fam_noL_* contains two non-learning or static models, namely “change detection” (CD) and “linear change detection” (LinCD). Both assume that the brain simply compares each incoming sensation to the preceding one. In model CD, *X_k_* at trial *k* is assigned to 0 if the *k^th^* stimulus is equal to the preceding one, and 1 otherwise. Model LinDC is similar to CD but assigns a prediction error proportional to the number of preceding sounds that differ from the one being currently observed (Lieder et al., 2013; Ostwald et al., 2012).
- Family *fam_L_* includes Bayesian perceptual learning models assuming that the brain estimates the probability *μ* to hear a standard (under a Bernoulli distribution, see Methods). These models involve a temporal integration window that allows down-weighting the influence of past events and whose size is defined by a time constant parameter (*τ*). We considered four different values for *τ* (2, 6, 10 and 100), leading to four models in *fam_L_*. precision-weighted prediction error computed at each trial quantifies the belief update about *μ* and was defined as the Bayesian Surprise (BS; the Kullback-Leibler divergence between the prior and the posterior distribution of *μ*).

These models were all fitted to the reconstructed cortical activity in both PC and UC conditions. For each source and at each time sample of the peristimulus interval, model inversions were performed following a meta-Bayesian approach using the VBA toolbox (Daunizeau et al., 2014) and model families were compared to each other using an RFX family-level inference (Penny et al., 2010).

#### 2. Results

As shown in Figure A1.B, the Null hypothesis (*famnull*) was found to outperform the other families, at every time sample and every cortical source, except for the time interval of the MMN. Precisely, posterior exceedance probability of family *fam_L_* was found to be significantly greater than the ones of the other families between 150 ms and 200 ms (6 samples), in the left and right Heschl’s Gyrus, between 130 ms and 180 ms (6 samples) and at 150 ms (1 sample) in the left and right Planum Polare, respectively, and between 130 ms and 200 ms (8 samples) and between 140 ms and 190 ms (6 samples) in the left and right Inferior Frontal Gyrus, respectively. Noticeably, in the latency range of the P3a described in Lecaignard et al.(2015), the Null hypothesis could also be challenged by learning models, as suggested by the increase of *fam_L_* posterior exceedance probability, an effect most visible in left IFG.

### A2. DCM analysis of surprise processing

We here address the neurophysiological mechanisms underlying perceptual learning during oddball processing. Using DCM, we test the hypothesis that precision-weighted prediction errors are conveyed upwards in the cortical hierarchy through feedforward connections, and trigger descending prediction messages. DCMs for evoked responses rely on a mesoscopic model of the cortical laminar circuitry (*Canonical Micro-Circuit*, CMC, Figure S2) which enables testing formally the mapping of computational learning quantities onto cortical representations (Bastos et al., 2012; Moran et al., 2013). Inspired by previous DCMs of the MMN (Auksztulewicz and Friston, 2016; Garrido et al., 2009a), and guided by current learning model predictions, we addressed the two following questions: what is the structure of the network engaged in typical oddball sequence processingWithin this auditory network, what modulation of effective connectivity supports deviant compared to standard tone processing? Each is addressed using Bayesian model comparison performed over a dedicated model space. This twofold analysis combines in a Bayesian way posterior DCMs inferred separately in conditions UC and PC, using EEG and MEG signals (details about the posterior fusion of EEG and MEG DCMs, denoted p-MEEG, is provided in the main manuscript).

#### 1. Methods

##### Model space for network characterization

We start by characterizing network architecture and input. Most reports of the DCM of the MMN entailed a three-level hierarchy (Auksztulewicz and Friston, 2015; Chennu et al., 2016; Garrido et al., 2009b; 2018; Phillips et al., 2015). We considered the six above-mentioned sources of the MMN. However, the complementary EEG and MEG topographical information regarding the predictability modulation of the MMN peak led us to test an additional level. Indeed, this predictability effect was larger at fronto-central sites in EEG (Lecaignard et al., 2015) and at temporal gradiometers in MEG (Figure 1.B), in a way that suggests superior frontal generators expressing poorly on frontal gradiometers. We could identify bilateral clusters of activity in this region over the significant time intervals reported in Lecaignard et al. (2015), with however limited precision. In short, left and right frontal clusters (36 and 29 nodes respectively, thresholded at p < 0.05 with Family Wise Error (FWE) whole-brain correction) were found for the early deviance effect, and a left contribution of 34 nodes for the MMN interval (with p < 0.001 not corrected). Each of the 8 resulting clusters led to an equivalent current dipole (ECD) located at the averaged position of local maxima over the different time intervals. MNI coordinates are provided in Figure A3.A. The resulting four-level DCM structure was composed of eight sources distributed bilaterally over (from the lowest to the highest level) HG, PP, IFG and SF. We connected these sources with extrinsic (forward and backward) connections. Alternative hypotheses entailed two- and three-level networks allowing to test the hierarchical depth as well as the contribution of PP and SF sources, leading to five model families (*A1*, *A2*, *A3*, *A4* and *A5;* see Figure A2). Regarding DCM inputs, all models included a direct input to bilateral HG. In addition, inputs targeting IFG sources (known to receive direct thalamic afferents) were tested as the source-reconstructed EEG and MEG evoked responses suggested that frontal regions were activated prior to temporal ones (Deouell, 2007). The input factor thereby included two levels (*HG* and *HG-IFG*). Importantly, the present model space does not address the presence or absence of trial-specific modulations (standard-to-deviant changes in connection strength) applying to extrinsic and intrinsic connections; this aspect is treated in the following. Of all intrinsic connections, we are concerned with the self-inhibition connection strength of the superficial pyramidal cells (subpopulation SP), hypothesized to encode precision (Bastos et al., 2012). Here, we assumed forward and backward trial-specific modulations, as already reported in several MMN-DCM studies (Garrido et al., 2009b), and we integrated over the two possible hypotheses (presence or absence) for the intrinsic modulation. DCM with CMC also includes extrinsic *modulatory* connections to enable the top-down indexation of subpopulation SP excitability on the output activity of higher-level feedbacking sources. We integrate over the two alternative hypotheses (presence or absence). The resulting model space thus comprised a total of 36 DCMs (Figure A2).

**Figure A2.**
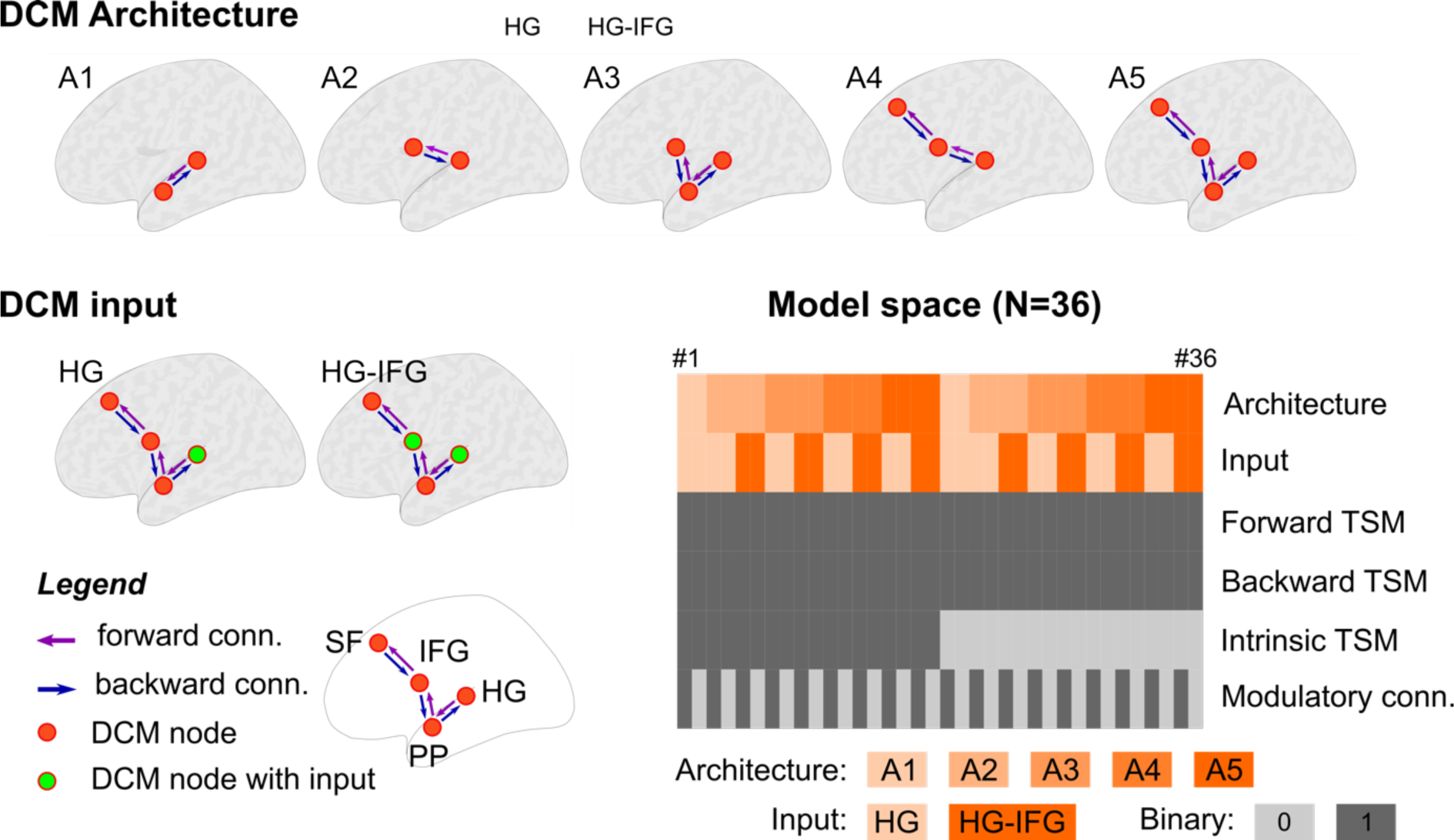
DCM model space (network structure analysis). Upper row, schematic view of the five model families designed to test DCM architecture in deviance processing. Bottom left, the two model families of DCM input, HG and HG-IFG. Color codes of extrinsic connections (conn.) and DCM source (or node) are provided in the legend. Source labels HG, PP, IFG and SF: Heschl’s Gyrus, Planum Polare, Inferior Frontal Gyrus and Superior Frontal. Bottom Right Table: DCM specifications for each of the 36 models (in columns). TSM: Trial-Specific Modulations accounting for standard-to-deviant changes in connectivity. Frontal, Backward and Intrinsic TSM, as well as Modulatory connections correspond to binary options (enabled = 1, disabled= 0) applying to the entire network.

**Figure A3.**
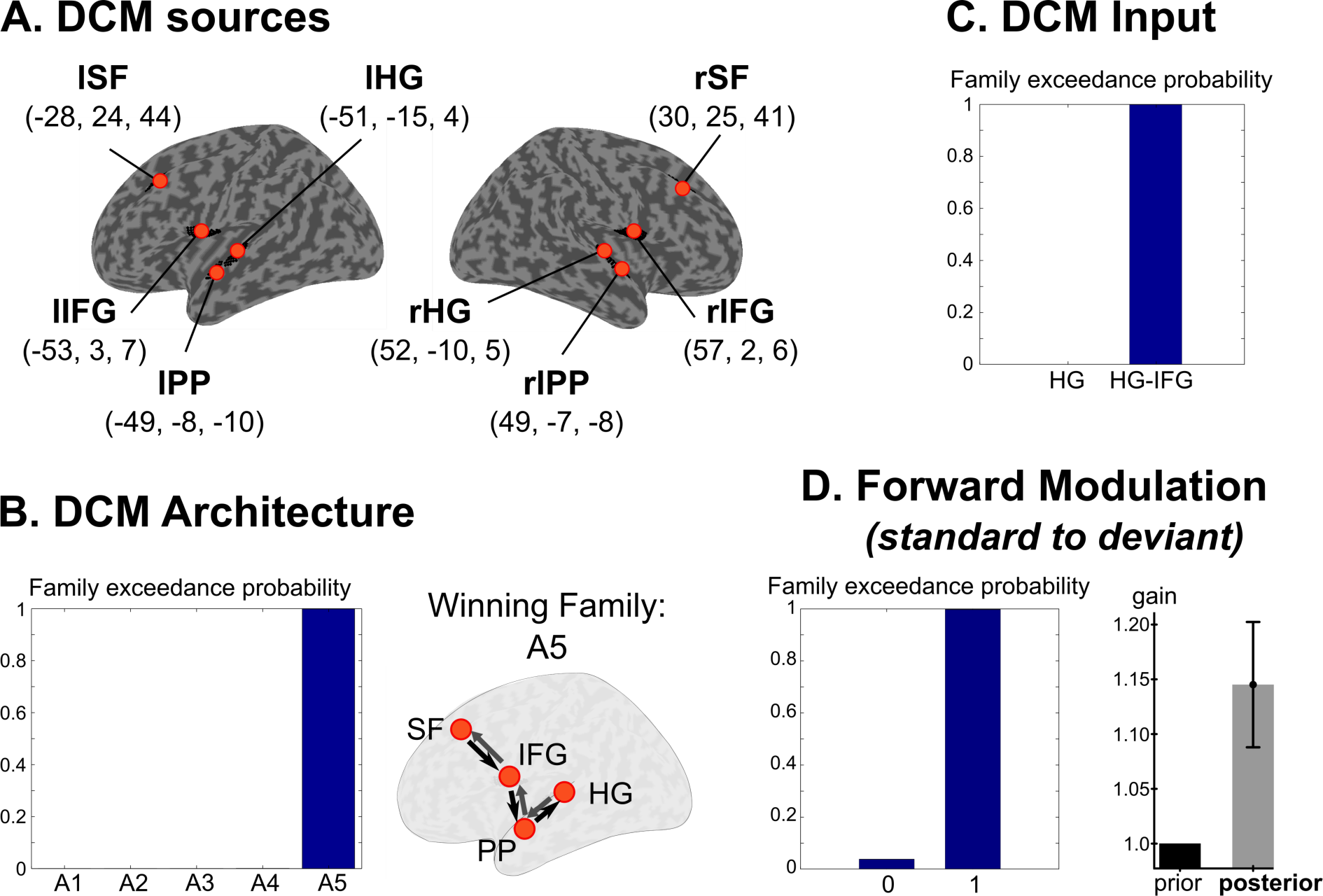
A. Dynamic causal modelling of deviance processing, using p-MEEG. A) Cortical sources for DCM analysis. Each source is indicated schematically with orange dots on the inflated cortex (SPM template brain), with corresponding MNI coordinates (mm) in parenthesis. B) Family inference for DCM architecture: family exceedance probabilities (left) and corresponding network for the winning family A5 (right) following the notations and color codes of Figure A2. C) Family inference for DCM inputs (family exceedance probabilities). D) Family inference for standard-to-deviant trial-specific modulation for the forward connectivity. Family exceedance probabilities of the model families with disabled (0) and enabled modulation (1) is provided (left) and group-level BMA estimates of gain values (in condition UC) averaged over the DCM network is represented (right). Prior value of gain (black bar) was set to 1 assuming no standard-to-deviant modulation. Labels for the cortical sources and model families are provided in the main text.

##### Model space for deviance-related changes in connectivity

This second model space aims at finessing the description of the extrinsic and the intrinsic connectivity within the winning DCM structure (architecture *A5* and double-input *HG-IFG*). We addressed in particular trial-specific gain applying on forward, backward, and intrinsic connections. For forward and intrinsic gains, two model families were considered each, (present or absent at all or none connections). Regarding backward gains, their effect depends on the above-mentioned extrinsic modulatory connections, if any. Specifically, they apply to backward extrinsic connections in the absence of modulatory connections, and to intrinsic connections otherwise (Bastos et al., 2012). This led us to consider three model families corresponding to backward gain i) disabled, ii) enabled without and iii) enabled with modulatory connections. A total of 14 models composed this model space.

##### Model inversion

DCMs were inverted with SPM12 (Wellcome Department of Imaging Neuroscience, http://www.fil.ion.ucl.ac.ik/spm). We used default values of SPM12 as prior expectations and prior variance for each DCM parameter to be estimated. Each DCM inversion involved standard response as the initial state of the system and deviant response resulting from the experimental perturbation. We infer the synaptic connection strengths as well as their trial-specific changes, and DCM sources orientation (but not their position). For each modality (EEG, MEG), the forward model used for DCM inversion was an advanced realistic Boundary Element Model (BEM) computed with Openmeeg software (Gramfort et al., 2010). Each model inversion was performed in condition UC and PC and for EEG and MEG data separately (leading to four inversions per subject and per model).

##### Statistical analysis

In the first analysis with the *network characterization* model space, we used data in condition UC and PC. For each condition, DCMs obtained in each modality (EEG, MEG) were fused using the above-described approach. We then combined p-MEEG DCMs across conditions (UC, PC) using similar Bayesian reasoning (log-posterior model evidences were summed across conditions). We then quantitatively evaluated the architecture and the input families using family-level inference (Penny et al., 2010) with an RFX model, based on the p-MEEG approximations of model evidence (ℱ*_p-MEEG_*). The second analysis with the deviance-*related changes* model space rests on p-MEEG DCMs in conditions UC and PC. Family level inference (RFX model) combined both conditions, and was conducted over the forward, backward and intrinsic gain parameters. As each of these three family comparisons indicated standard-to-deviant changes, we subsequently examined gain estimates in each condition separately to determine the direction of change (a value larger than one would indicate larger connection strength in deviants than in standards, and *vice-versa*). For each condition, we used Bayesian model averaging (BMA) to derive group-level posterior estimates averaged across model space (with model-evidence weighting) and across subjects. Each estimate was informed by EEG and MEG using equation Eq.11. For each connection type, BMA estimates were averaged over the entire network.

#### 2. Results

##### Network characterization

Resulting explained variance averaged across models and subjects was equal 92.5% (standard deviation: ±10.5) and 78.1% (±11.6) in EEG and MEG, respectively (condition UC), and to 91.9% (±12.1) and 78.1% (±11.6) in EEG and MEG, resp. (condition PC). In the p-MEEG modality, family level inference revealed that family *A5* outperformed other model families with a posterior confidence probability (*pcp*) and posterior exceedance probability (*pep*) of 0.68 and >0.99, respectively (Figure A3.B). Regarding the DCM inputs (Figure A3.C), family level inference was clearly in favor of models with inputs arriving in both HG and IFG sources (*pcp* / *pep*: 0.82 / >0.99).

##### Deviance-related changes in connectivity

Explained variance was equal to (results for condition UC / condition PC): 96.5% (±5.1) / 97.8% (±2.7), for EEG, and 87.1% (±7.4) / 87.5% (±5.5) for MEG. We found evidence for forward, backward and intrinsic deviant modulation (*pcp* / *pep*: 0.82 / >0.99; 0.73 / 0.99; 0.73 / 0.99, resp.) in line with pioneering DCMs of the MMN (Auksztulewicz and Friston, 2016; Garrido et al., 2009b). In order to further characterize the deviant-related direction of change, we examined group-level posterior estimates of trial-specific gain obtained with Bayesian model averaging (BMA) over model space and subjects. Over the entire network and in both conditions, we found standard-to-deviant modulations that fits well with predictive coding message-passing expectations. In particular, we found larger forward coupling for deviants (Figure A3.D; average and standard error across the network of group-level BMA estimates of forward trial-specific gain. UC: 1.145 ± 0.057, present in 5 out of 6 forward connections; PC: 1.027 ± 0.042, 4 out of 6 connections). We also found the expected increase of backward gain (UC: 1.179 ± 0.073, 5 out of 6 connections; PC: 1.307 ± 0.112, 5 out of 6 connections) and a decrease of the intrinsic gain (UC: 0.947 ± 0.064, 5 out of 8 connections. PC: 0.956 ± 0.047, 5 out of 8 connections).

---

**Figure S1.**
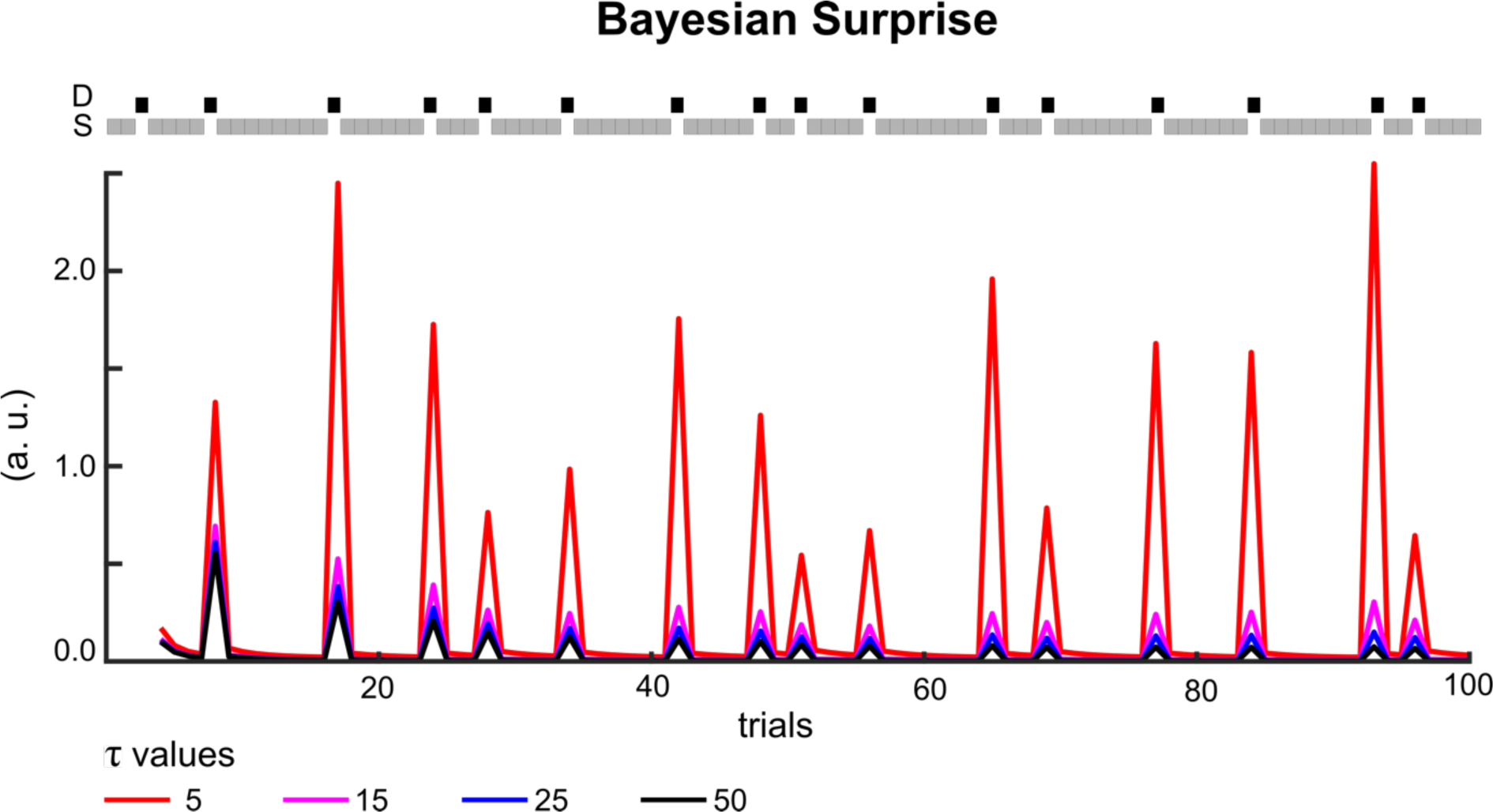
Bayesian surprise as a function of *τ*. (arbitrary units, a. u.) Illustration of different BS trajectories obtained with varying *τ*, for the first 100 stimuli of a typical UC oddball sequence. Two comments should be made: 1) BS decreases as *τ* increases and 2) whatever *τ*, BS is larger for deviants (D, black squares) than for standards (S, grey squares).

**Figure S2.**
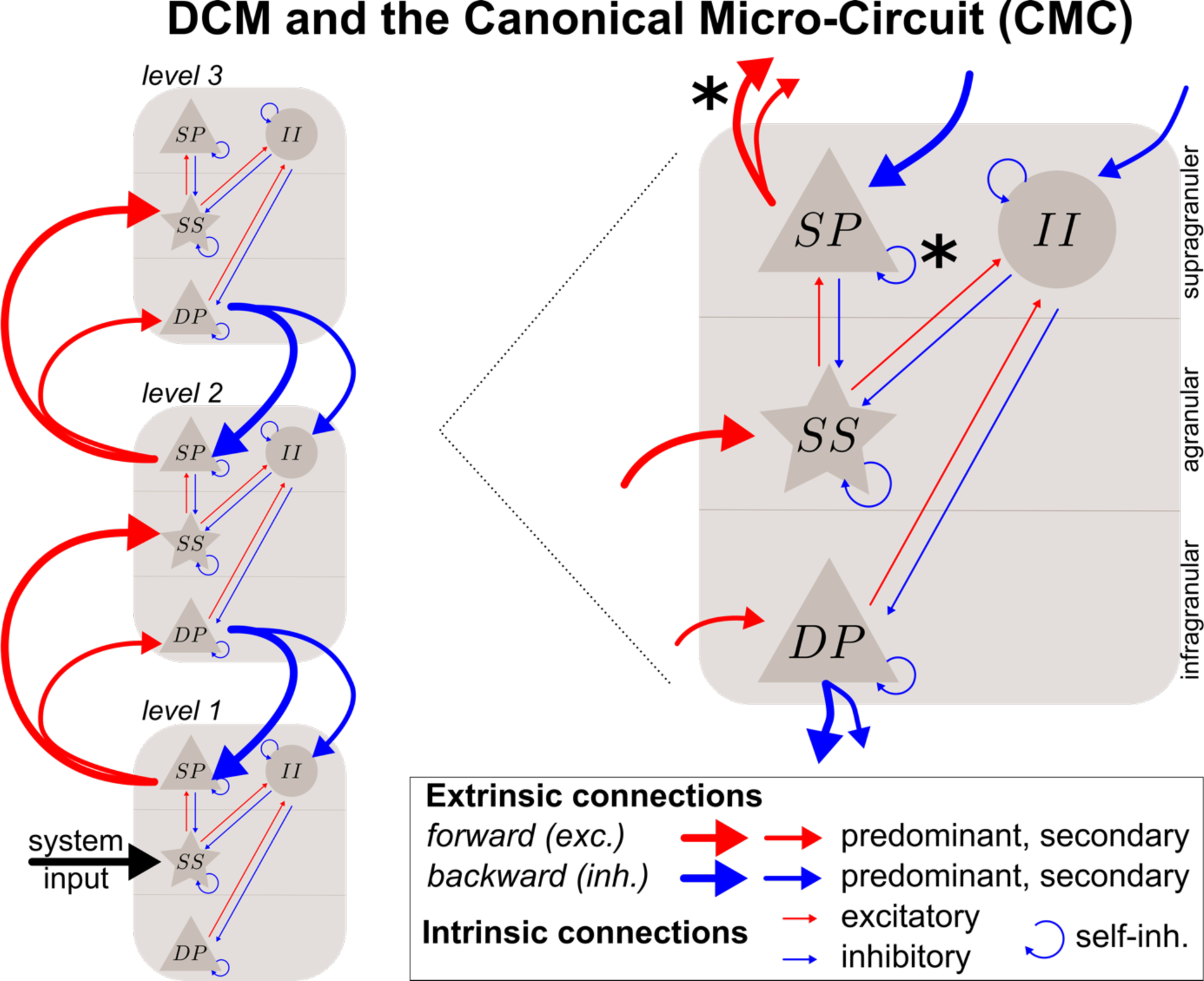
Schematic description of DCM with the Canonical Micro-Circuit. Left, example of a three-level DCM with sensory input (black arrow) targeting the lower level. Color and strength of connections indicate their category, as indicated in the legend. Right, detailed view of a DCM source (level), embedding a canonical micro-circuitry. It comprises three layers (infragranular, agranular and supragranular) and involves four cell suppopulations (DP: deep pyramidal; SS: spiny stellate; SP: superior pyramidal; II: inhibitory interneurons). Black stars indicate the predominant forward and self-inhibitory connections that were examined in the DCM-predictability analysis, because they could represent prediction error (or precision-weighted prediction error) and precision, respectively, according to predictive brain models. Trial-specific modulations reflecting standard-to-deviant changes are not represented. Forward and backward modulations apply on forward and backward extrinsic connections, respectively, and intrinsic one applies on self-inhibition in SP. exc.: excitatory; inh.: inhibitory.

**Table S1.** Sufficient statistics (mean, variance) for initial conditions and model parameters. Ev. Par.: Evolution parameters; Obs. Par.: Observation parameters. For family fam_noL_, previous observation set to 0 correspond to a standard sound. *θ*_1_ and *θ*_2_) refers to Equation Eq.1. Initial states values were not estimated during model inversion (variance is set to 0).

| family | model | Initial state values |  | Ev. Par |  | Obs. Par |  |
| --- | --- | --- | --- | --- | --- | --- | --- |
| fam<sub>null</sub> | null | - | - | - | - | $\theta_2$ | (0, 100) |
| fam<sub>noL</sub> | CD | previous observation | (0, 0) | - | - | $\theta_1$ | (0, 100) |
| | | | | | | $\theta_2$ | (0, 100) |
| | LinCD | previous observation | (0, 0) | - | - | $\theta_1$ | (0, 100) |
| | | observed standards | (0, 0) | - | - | $\theta_2$ | (0, 100) |
| fam<sub>L</sub> | fixed $\tau$ | standard count | (1, 0) | - | - | $\theta_1$ | (0, 100) |
| | | deviant count | (1, 0) | - | - | $\theta_2$ | (0, 100) |
| | free $\tau$ | standard count | (1, 0) | $\tau$ | (log(10), 10) | $\theta_1$ | (0, 100) |
| | | deviant count | (1, 0) | | | $\theta_2$ | (0, 100) |

## Notes

### Competing Interest Statement

The authors have declared no competing interest.

## References

Adams, R.A., Stephan, K.E., Brown, H.R., Frith, C.D., Friston, K., 2013. The computational anatomy of psychosis. Front Psychiatry 4, 47. doi:10.3389/fpsyt.2013.00047

Aguera, P.-E., Jerbi, K., Caclin, A., Bertrand, O., 2011. ELAN: A Software Package for Analysis and Visualization of MEG, EEG, and LFP Signals. Computational Intelligence and Neuroscience 2011, 1–11. doi:10.1155/2011/158970

Auksztulewicz, R., Barascud, N., Cooray, G.K., Nobre, A.C., Chait, M., Friston, K., 2017. The Cumulative Effects of Predictability on Synaptic Gain in the Auditory Processing Stream. Journal of Neuroscience 37, 6751–6760. doi:10.1523/JNEUROSCI.0291-17.2017

Auksztulewicz, R., Friston, K., 2016. Repetition suppression and its contextual determinants in predictive coding. CORTEX. doi:10.1016/j.cortex.2015.11.024

Auksztulewicz, R., Friston, K., 2015. Attentional Enhancement of Auditory Mismatch Responses: a DCM/MEG Study. Cerebral Cortex. doi:10.1093/cercor/bhu323

Bastos, A.M., Usrey, W.M., Adams, R.A., Mangun, G.R., Fries, P., Friston, K., 2012. Canonical microcircuits for predictive coding. Neuron 76, 695–711. doi:10.1016/j.neuron.2012.10.038

Brown, H.R., Friston, K., 2013. The functional anatomy of attention: a DCM study. Front Hum Neurosci 7, 784. doi:10.3389/fnhum.2013.00784

Brown, H.R., Friston, K., 2012. Dynamic causal modelling of precision and synaptic gain in visual perception - an EEG study. NeuroImage 63, 223–231. doi:10.1016/j.neuroimage.2012.06.044

Chennu, S., Noreika, V., Gueorguiev, D., Blenkmann, A., Kochen, S., Ibáñez, A., Owen, A.M., Bekinschtein, T.A., 2013. Expectation and attention in hierarchical auditory prediction. Journal of Neuroscience 33, 11194–11205. doi:10.1523/JNEUROSCI.0114-13.2013

Chennu, S., Noreika, V., Gueorguiev, D., Shtyrov, Y., Bekinschtein, T.A., Henson, R., 2016. Silent Expectations: Dynamic Causal Modeling of Cortical Prediction and Attention to Sounds That Weren’t. J Neurosci 36, 8305–8316. doi:10.1523/JNEUROSCI.1125-16.2016

Clark, A., 2013. Whatever next? Predictive brains, situated agents, and the future of cognitive science. Behav Brain Sci 36, 181–204. doi:10.1017/S0140525X12000477

Daunizeau, J., Adam, V., Rigoux, L., 2014. VBA: a probabilistic treatment of nonlinear models for neurobiological and behavioural data. PLoS Comput Biol 10, e1003441. doi:10.1371/journal.pcbi.1003441

Dayan, P., Hinton, G.E., Neal, R.M., Zemel, R.S., 1995. The Helmholtz machine. Neural Comput 7, 889–904.

Feldman, H., Friston, K., 2010. Attention, uncertainty, and free-energy. Front Hum Neurosci 4, 215. doi:10.3389/fnhum.2010.00215

Fitzgerald, K., Todd, J., 2020. Making Sense of Mismatch Negativity. Front Psychiatry 11, 468. doi:10.3389/fpsyt.2020.00468

Fogelson, N., Litvak, V., Peled, A., Fernandez-del-Olmo, M., Friston, K., 2014. The functional anatomy of schizophrenia: A dynamic causal modeling study of predictive coding. Schizophr Res 158, 204–212. doi:10.1016/j.schres.2014.06.011

Friston, K., 2020. Bayesian Dysconnections. Am J Psychiatry 177, 1110–1112. doi:10.1176/appi.ajp.2020.20091421

Friston, K., 2012. The history of the future of the Bayesian brain. NeuroImage 62, 1230–1233. doi:10.1016/j.neuroimage.2011.10.004

Friston, K., 2010. The free-energy principle: a unified brain theory? Nat Rev Neurosci 11, 127–138. doi:10.1038/nrn2787

Friston, K., 2008a. Hierarchical models in the brain. PLoS Comput Biol 4, e1000211. doi:10.1371/journal.pcbi.1000211

Friston, K., 2008b. Hierarchical Models in the Brain 4, e1000211. doi:10.1371/journal.pcbi.1000211

Friston, K., 2005. A theory of cortical responses. Philosophical Transactions of the Royal Society B: Biological Sciences 360, 815–836. doi:10.1098/rstb.2005.1622

Friston, K., Harrison, L.M., Daunizeau, J., Kiebel, S., Phillips, C., Trujillo-Barreto, N.J., Henson, R., Flandin, G., Mattout, J., 2008. Multiple sparse priors for the M/EEG inverse problem. NeuroImage 39, 1104–1120. doi:10.1016/j.neuroimage.2007.09.048

Friston, K., Kiebel, S., 2009. Predictive coding under the free-energy principle. Journal of Mathematical Psychology 364, 1211–1221. doi:10.1098/rstb.2008.0300

Garrido, M.I., Kilner, J., Kiebel, S., Friston, K., 2009. Dynamic causal modeling of the response to frequency deviants. Journal of Neurophysiology 101, 2620–2631. doi:10.1152/jn.90291.2008

Gramfort, A., Papadopoulo, T., Olivi, E., Clerc, M., 2010. OpenMEEG: opensource software for quasistatic bioelectromagnetics. Biomed Eng Online 9, 45. doi:10.1186/1475-925X-9-45

Haarsma, J., Kok, P., Browning, M., 2020. The promise of layer-specific neuroimaging for testing predictive coding theories of psychosis. Schizophr Res 1–9. doi:10.1016/j.schres.2020.10.009

Heilbron, M., Chait, M., 2018. Great Expectations: Is there Evidence for Predictive Coding in Auditory Cortex? Neuroscience 389, 54–73. doi:10.1016/j.neuroscience.2017.07.061

Henson, R., Mouchlianitis, E., Friston, K., 2009. MEG and EEG data fusion: simultaneous localisation of face-evoked responses. NeuroImage 47, 581–589. doi:10.1016/j.neuroimage.2009.04.063

Kanai, R., Komura, Y., Shipp, S., Friston, K., 2015. Cerebral hierarchies: predictive processing, precision and the pulvinar. Philosophical Transactions of the Royal Society B: Biological Sciences 370, 20140169–20140169. doi:10.1098/rstb.2014.0169

Kiebel, S., Garrido, M.I., Moran, R., Chen, C.C., Friston, K., 2009. Dynamic causal modeling for EEG and MEG. Hum. Brain Mapp. 30, 1866–1876. doi:10.1002/hbm.20775

Lawson, R.P., Mathys, C.D., Rees, G., 2017. Adults with autism overestimate the volatility of the sensory environment. Nat Neurosci 15, 173–13. doi:10.1038/nn.4615

Lecaignard, F., Bertrand, O., Caclin, A., Mattout, J., 2021. Empirical Bayes evaluation of fused EEG-MEG source reconstruction: Application to auditory mismatch evoked responses. NeuroImage 226, 117468. doi:10.1016/j.neuroimage.2020.117468

Lecaignard, F., Bertrand, O., Gimenez, G., Mattout, J., Caclin, A., 2015. Implicit learning of predictable sound sequences modulates human brain responses at different levels of the auditory hierarchy. Front Hum Neurosci 9, 505. doi:10.3389/fnhum.2015.00505

Lieder, F., Daunizeau, J., Garrido, M.I., Friston, K., Stephan, K.E., 2013. Modelling trial-by-trial changes in the mismatch negativity. PLoS Comput Biol 9, 1–16. doi:10.1371/journal.pcbi.1002911

Litvak, V., Friston, K., 2008. Electromagnetic source reconstruction for group studies. NeuroImage 42, 1490–1498. doi:10.1016/j.neuroimage.2008.06.022

Lopes da Silva, F., 2013. EEG and MEG: relevance to neuroscience. Neuron 80, 1112–1128. doi:10.1016/j.neuron.2013.10.017

Lumaca, M., Dietz, M.J., Hansen, N.C., Quiroga-Martinez, D.R., Vuust, P., 2021. Perceptual learning of tone patterns changes the effective connectivity between Heschl’s gyrus and planum temporale. Hum. Brain Mapp. 42, 941–952. doi:10.1002/hbm.25269

Lumaca, M., Trusbak Haumann, N., Brattico, E., Grube, M., Vuust, P., 2018. Weighting of neural prediction error by rhythmic complexity: A predictive coding account using mismatch negativity. Eur J Neurosci. doi:10.1111/ejn.14329

Mathys, C.D., Lomakina, E.I., Daunizeau, J., Iglesias, S., Brodersen, K.H., Friston, K., Stephan, K.E., 2014. Uncertainty in perception and the Hierarchical Gaussian Filter. Front Hum Neurosci 8, 825. doi:10.3389/fnhum.2014.00825

Mattout, J., Phillips, C., Penny, W.D., Rugg, M.D., Friston, K., 2006. MEG source localization under multiple constraints: an extended Bayesian framework. NeuroImage 30, 753–767. doi:10.1016/j.neuroimage.2005.10.037

Meyniel, F., 2020. Brain dynamics for confidence-weighted learning. PLoS Comput Biol 16, e1007935. doi:10.1371/journal.pcbi.1007935

Meyniel, F., Maheu, M., Dehaene, S., 2016. Human Inferences about Sequences: A Minimal Transition Probability Model. PLoS Comput Biol 12, e1005260. doi:10.1371/journal.pcbi.1005260

Moran, R., Campo, P., Symmonds, M., Stephan, K.E., Dolan, R.J., Friston, K., 2013. Free energy, precision and learning: the role of cholinergic neuromodulation. Journal of Neuroscience 33, 8227–8236. doi:10.1523/JNEUROSCI.4255-12.2013

Ostwald, D., Spitzer, B., Guggenmos, M., Schmidt, T.T., Kiebel, S., Blankenburg, F., 2012. Evidence for neural encoding of Bayesian surprise in human somatosensation. NeuroImage 62, 177–188. doi:10.1016/j.neuroimage.2012.04.050

Parr, T., Friston, K., 2018. Attention or salience? Curr Opin Psychol 29, 1–5. doi:10.1016/j.copsyc.2018.10.006

Penny, W.D., Mattout, J., Trujillo-Barreto, N.J., 2006. Bayesian model selection and averaging, in: Statistical Parametric Mapping the Analysis of Functional Brain Images. Elsevier, London, pp. 1–29.

Phillips, H.N., Blenkmann, A., Hughes, L.E., Kochen, S., Bekinschtein, T.A., Cam- CAN, Rowe, J.B., 2016. Convergent evidence for hierarchical prediction networks from human electrocorticography and magnetoencephalography. CORTEX 82, 192–205. doi:10.1016/j.cortex.2016.05.001

Pinotsis, D.A., Geerts, J.P., Pinto, L., Fitzgerald, T.H.B., Litvak, V., Auksztulewicz, R., Friston, K., 2017. Linking canonical microcircuits and neuronal activity_ Dynamic causal modelling of laminar recordings. NeuroImage 146, 355–366. doi:10.1016/j.neuroimage.2016.11.041

Rohenkohl, G., Cravo, A.M., Wyart, V., Nobre, A.C., 2012. Temporal expectation improves the quality of sensory information. Journal of Neuroscience 32, 8424–8428. doi:10.1523/JNEUROSCI.0804-12.2012

Schröger, E., Kotz, S.A., Sanmiguel, I., 2015. Bridging prediction and attention in current research on perception and action. Brain Res 1626, 1–13. doi:10.1016/j.brainres.2015.08.037

Southwell, R., Baumann, A., Gal, C., Barascud, N., Friston, K., Chait, M., 2017. Is predictability salient? A study of attentional capture by auditory patterns. Journal of Mathematical Psychology 372. doi:10.1098/rstb.2016.0105

Spratling, M.W., 2016. A review of predictive coding algorithms. Brain Cogn. doi:10.1016/j.bandc.2015.11.003

Stefanics, G., Heinzle, J., Horváth, A.A., Stephan, K.E., 2018. Visual Mismatch and Predictive Coding: A Computational Single-Trial ERP Study. Journal of Neuroscience 38, 4020–4030. doi:10.1523/JNEUROSCI.3365-17.2018

Tillmann, B., Bigand, E., Pineau, M., 1998. Effects of Global and Local Contexts on Harmonic Expectancy. Music Perception: An Interdisciplinary Journal 16, 99–117. doi:10.2307/40285780

Walsh, K.S., McGovern, D.P., Clark, A., O’Connell, R.G., 2020. Evaluating the neurophysiological evidence for predictive processing as a model of perception. Ann. N. Y. Acad. Sci. 1464, 242–268. doi:10.1111/nyas.14321

Weber, L.A.E., Diaconescu, A.O., Mathys, C.D., Schmidt, A., Kometer, M., Vollenweider, F., Stephan, K.E., 2020. Ketamine Affects Prediction Errors about Statistical Regularities: A Computational Single-Trial Analysis of the Mismatch Negativity. Journal of Neuroscience 40, 5658–5668. doi:10.1523/JNEUROSCI.3069-19.2020

## References

Deouell, L.Y., 2007. The Frontal Generator of the Mismatch Negativity Revisited. Journal of Psychophysiology 21, 188–203. doi:10.1027/0269-8803.21.3.188

Garrido, M.I., Kilner, J., Kiebel, S., Friston, K., 2009a. Dynamic causal modeling of the response to frequency deviants. Journal of Neurophysiology 101, 2620–2631. doi:10.1152/jn.90291.2008

Garrido, M.I., Kilner, J., Stephan, K.E., Friston, K., 2009b. The mismatch negativity: a review of underlying mechanisms. Clin Neurophysiol 120, 453–463. doi:10.1016/j.clinph.2008.11.029

Garrido, M.I., Rowe, E.G., Halasz, V., Mattingley, J.B., 2018. Bayesian Mapping Reveals That Attention Boosts Neural Responses to Predicted and Unpredicted Stimuli. Cereb Cortex 28, 1771–1782. doi:10.1093/cercor/bhx087

Mattout, J., Henson, R., Friston, K., 2007. Canonical source reconstruction for MEG. Computational Intelligence and Neuroscience 67613. doi:10.1155/2007/67613

Penny, W.D., Stephan, K.E., Daunizeau, J., Rosa, M.J., Friston, K., Schofield, T.M., Leff, A.P., 2010. Comparing families of dynamic causal models. PLoS Comput Biol 6, e1000709. doi:10.1371/journal.pcbi.1000709

Phillips, H.N., Blenkmann, A., Hughes, L.E., Bekinschtein, T.A., Rowe, J.B., 2015. Hierarchical Organization of Frontotemporal Networks for the Prediction of Stimuli across Multiple Dimensions. Journal of Neuroscience 35, 9255–9264. doi:10.1523/JNEUROSCI.5095-14.2015

